# Hand preference predicts behavioral responses to threats in Barbary macaques

**DOI:** 10.1101/2023.01.06.523039

**Authors:** Eva S.J. van Dijk, Debottam Bhattacharjee, Elena Belli, Jorg J.M. Massen

**Affiliations:** Animal Behavior and Cognition, Department of Biology, Utrecht University, Padualaan 8, 3584 CH Utrecht, The Netherlands; IFM Biology, Department of Physics, Chemistry and Biology, Linköping University, Olaus Magnus väg 37, 583 30 Linköping, Sweden

**Keywords:** brain lateralization, handedness, non-human primates, individual variation, anti-predator response

## Abstract

The structure and functioning of the brain are lateralized – the right hemisphere processes unexpected stimuli and controls spontaneous behavior, while the left deals with familiar stimuli and routine responses. Hemispheric dominance, the predisposition of an individual using one hemisphere over the other, may lead to behavioral differences; particularly, an individual may be programmed to act in a certain way concerning hemispheric dominance. Hand preference is a robust estimator of hemispheric dominance in primates, as each brain hemisphere controls the opposing side of the body. Studies have found links between hand preference and the exhibition of different behaviors in contexts such as exploring and manipulating objects. However, little is known about whether hand preference can predict behavioral variations in other ecologically relevant contexts like, for example, predation. We investigated the relationship between hand preference and the behavioral responses to two types of predator models in captive Barbary macaques (*Macaca sylvanus*) (n=22). Hand preference was determined by observing unimanual foraging, whereas focus and tension behaviors were quantified during experimental exposure to predator models. We found 91% of the macaques to be lateralized with no group-level bias. In contrast to their right-hand counterparts, individuals with a strong left-hand preference elicited frequent focus and tension behavior. Additionally, the behavioral response varied with predator type. We also found an interaction effect between hand preference and the predator type. Our study suggests that hand preference can reliably predict behavioral variations in the context of potential predation. While these results are consistent with the lateralized brain function, indicating lateralization as a potential neural mechanism of behavioral variation, the interaction effect between hand preference and predator type elucidates the importance of context-specificity when investigating laterality non-invasively. Future research on other non-human primates using the current framework may shed light on the evolution of laterality and underlying behavioral predispositions.

## 1. Introduction

The left and right hemispheres control opposing sides of the body, process information differently, and control contrasting behaviors. Such asymmetries in the structure and/or function of the two hemispheres are called cerebral lateralization (Bisazza et al., 1998). The contrasting behavioral responses linked to cerebral lateralization follow the same basic pattern among vertebrates (Rogers, 2002; Rogers & Andrew, 2002). Roughly, the left hemisphere controls routine behavior and functions, while the right one detects and responds to novel and unexpected stimuli like responses to potential predators. Overall, the left hemisphere is said to be involved when controlling responses that first require weighing different options, while the right hemisphere is involved in spontaneous and intense reactions to various stimuli (Rogers, 2002, 2010).

Next to the processing of stimuli and controlling behaviors, both hemispheres are involved in regulating short-term affective states, i.e., emotions (Désiré et al., 2002). Like behavioral responses, emotional functioning seems lateralized across human and non-human primates (Leliveld et al., 2013). The left hemisphere is dominant for positive emotions, at least concerning responses to food rewards; on the contrary, the right hemisphere is specialized in expressing intense, often negative emotions, such as fear and disgust, which likely relates to the specialization for the control of aggressive interactions (Rogers, 2002). Similarly, the right hemisphere is associated with the stress response (Ocklenburg et al., 2016; Rogers, 2010). For example, in black-tufted marmosets (*Callithrix penicillata*), there is a stronger right hemisphere activation under acute stress as measured by tympanic temperature (Tomaz et al., 2003); and in rhesus macaques (*Macaca mulatta*), plasma cortisol levels positively correlate with activity levels in the right frontal cortex, associated with high levels of fear and defensive behavior (Kalin et al., 1998). As each hemisphere processes stimuli differently and controls distinct sets of behaviors and emotions, hemispheric specialization may be considered a neurobiological mechanism underlying intraspecific behavioral variation (Rogers, 2009, 2010). However, non-invasive measurements are incredibly difficult to implement while investigating the role of cerebral lateralization as a predictor of behavioral variation.

Across cultures, by far, most humans are right-handed for manipulative actions (Raymond & Pontier, 2004), yet there exists a considerable variation in hand preference both within and across other primate species (Meguerditchian et al., 2013; Kasper et al., 2022; Soto et al., 2022). Each side of the body is controlled by the opposite brain hemisphere; i.e., the right hand is controlled by the left hemisphere and vice versa (Bisazza et al., 1998). When there is a group-level hand bias for a task, such as human right-hand bias for manual tasks, this could result from functional hemispheric specialization favoring the hemisphere best equipped for the task. However, when there is no group-level hand bias for a task and likely no functional hemispheric specialization, individual hand preference reflects an individual’s tendency to use the related hemisphere more than the other, a feature called hemisperic dominance (Hook & Rogers, 2000; Rogers, 2011). Hemispheric dominance may thus program an individual to act in a certain way, leading to behavioral variation.

Although bimanual tasks are proposed to be more suitable when investigating hand preference on a group level (Nelson, 2022; Soto et al., 2022), an individual’s hand preference for simple and cognitively less demanding tasks, such as unimanual reaching, also reflects one hemisphere’s dominance over the other (Gordon & Rogers, 2010, 2015; Rogers, 2018). Individual hand preference for unimanual reaching or unimanual foraging is highly consistent within individuals across primates (e.g., *Callithrix jacchus*: Kuběnová et al., 2022; *Macaca silenus*: Rogers, 2009; *Rhinopithecus roxellana*: Fu et al., 2022). Moreover, research on non-human primates found differences among right and left-handed individuals; the hand preference of common marmosets, for example, which is stable over adulthood and across tasks (Gordon & Rogers, 2015), also predicts the marmosets’ reaction to novelty, with left-handed individuals taking longer to enter novel rooms and touching or exploring novel objects (Cameron & Rogers, 1999). A different study also showed that left-handed marmosets were less responsive to their social group and less proactive when investigating novel stimuli (Gordon & Rogers, 2010). On the other hand, right-handed marmosets are said to be more explorative, inquisitive and proactive, and more influenced by and of influence to their social group (Rogers, 2018). A recent study, however, did not find a link between handedness and inter-individual behavioral differences, aka personality, in common marmosets (Masilkova et al., 2022). In Chimpanzees (*Pan troglodytes*), right-handed individuals appear more curious than left-handed individuals (Hopkins & Bennett, 1994). Similarly, in a comparative study on multiple primate species, left-handed individuals took longer before inspecting novel objects but were also less fearful and less inactive (Fernández-Lázaro et al., 2019). Overall, right-handed non-human primates are more likely to approach and interact with novel stimuli than their left-handed conspecifics, and studies in humans are consistent with this pattern (Rogers, 2018; Wright et al., 2013).

Non-human primate lateralization and hand preference in relation to behavior have been investigated in different contexts (Rogers, 2018). Yet, only a few studies looked at the effects of handedness on behavior in ecologically valid contexts. Zonato and colleagues (2022) argue that ecological factors should be evaluated when assessing hand preference. They tested hand preference in ring-railed lemurs (*Lemur catta*) and found evidence of individual hand preference for grasping static food items but not for food in motion, a dynamically complex condition. Examining ecological factors when evaluating hand preference is gaining importance but remains relatively understudied. Differences between left- and right-handed individuals have been investigated for context-specific behaviors such as exploration (Braccini & Caine, 2009; Fernández-Lázaro et al., 2019), social behavior (Westergaard et al., 2003), cognitive bias (Gordon & Rogers, 2015), fear and stress responses (Rogers, 2009) and learning (Cameron & Rogers, 1999), as well as performance in cognitive tasks (Wang et al., 2022). To validate and understand the implications of relationships between hand preference and behavioral measures, more ecologically valid measures are required. For example, little is known about whether hand preference predicts individual behavioral variation in contexts such as predation.

As predator avoidance is of high ecological relevance, it is interesting to investigate whether hemispheric specialization results in individual behavioral differences with regard to reactions to predators. Predation is considered a strong selective pressure driving primate evolution, including the evolution of sociality (Anderson, 1986; Kappeler & van Schaik, 2002). Encounters with both live and simulated predators can result in stress (Cheney & Seyfarth, 2009). To avoid being predated on, primates display antipredator behavior (Barros et al., 2008; Stanford, 2002), including elevated vigilance and avoidance strategies, such as alarm calls and flight, but sometimes also confrontation (i.e., mobbing). The failure to avoid predation may have severe survival consequences, including death; thus, appropriate antipredator responses are vital, yet, there are clear interindividual differences when it comes to responding to potential predators (Carter et al., 2012). For example, in common marmosets, inter-individual variation has been observed in response to potential predators, which is considered part of a non-social personality trait, ‘Boldness-Shyness in Predation’ (Šlipogor et al., 2016). The right hemisphere may be especially important to the behavioral responses to predators as the right hemisphere detects unexpected stimuli such as potential predators and controls spontaneous behaviors such as flight. Moreover, it is specialized in expressing negative emotions such as fear and controls the stress response. Individuals with a left-hand bias and thus right-hemispheric dominance might therefore be more reactive to predators. One of the few studies to date that we know of, which investigated the link between predator responses and handedness, indeed found that left-handed Geoffrey marmosets (*Callithrix geoffroyi*), when confronted with a hawk call, freeze for longer than right-handed individuals (Braccini & Caine, 2009). Conversly, when common marmosets are confronted with a threat, right-handed individuals produce more mobbing calls and perform more head cocking and paralax movements than left-handed individuals (Gordon & Rogers, 2010). Interindividual differences in hemispheric dominance could thus play a major role in the observed interindividual behavioral responses to predators.

Due to the high variation in the social, behavioral and ecological characteristics, macaques are considered highly relevant when investigating lateralization (Regaiolli et al., 2018). One species of interest for lateralization is the relatively socially tolerant Barbary macaque (*Macaca sylvanus*) (Thierry et al., 2000). To the best of our knowledge, only three studies have so far investigated hand preference in Barbary macaques. These studies reported seven out of twenty (35%), seven out of fifteen (47%), and nine out of twelve (75%) individuals to be significantly lateralized (Baldachini et al., 2021; Regaiolli et al., 2018; Schmitt et al., 2008). However, these studies used different methods to collect data on hand use and to determine hand preference. Nevertheless, the results of these studies showed that for sequences of unimanual reaching, older individuals had an overall stronger hand preference (Schmitt et al., 2008). They did not find any effect of dominance rank on hand preference during bouts of unimanual interactions with inanimate targets using both food and non-food (Baldachini et al., 2021). Finally, none of these studies reported group-level lateralization for unimanual reaching or manipulation of inanimate objects (Baldachini et al., 2021; Regaiolli et al., 2018; Schmitt et al., 2008). None of these studies, however, investigated the relationship between hand preference and response to predators in Barbary macaques.

The Barbary or North African leopard (*Panthera pardus panthera*) is expected to have been a major predator of the Barbary macaque when the two species were still sympatric (Bautista, 2019; Fooden, 2007). Currently, the main predator of Barbary macaques is the domestic dog (*Canis lupus familiaris*), but jackals (*Canis aureus*), genets (*Genetta genetta*) and some species of birds of prey are also suggested as potential predators (Bautista, 2019; Majolo et al., 2013; Waterman et al., 2020). Besides, Barbary macaques are known to respond to snakes. While fear of snakes is common among primates and catarrhines, Barbary macaques have also co-existed with venomous snakes throughout their evolutionary history (Isbell, 2006; Öhman & Mineka, 2003). The semi-free-ranging populations of these macaques living in monkey parks in Europe are known to give alarm calls to snakes (Fischer & Hammerschmidt, 2002). There are also anecdotal observations of Barbary macaques standing bipedally, peeking in surrounding grass, approaching and even mobbing a snake upon detection (Fischer & Hammerschmidt, 2001). Moreover, even the non-native population of Gibraltar responds to snakes, despite the present snake species posing no actual threat to infant macaques (Fooden, 2007; Roberts et al., 2008). Snakes are thus proposed to be an ecologically relevant stimulus for Barbary macaques in addition to felid predators.

While both felid predators and snakes may constitute a relevant threat to Barbary macaques, potentially eliciting antipredator behavior and stress, the response to each type of predator may differ. Antipredator behavior serves to avoid predation and can thus vary for different types of predators based on their hunting techniques, the perceived threat, the mode of detection and the habitat structure (Lemasson et al., 2009). For example, wild Campbell’s monkeys (*Cercopithecus c. campbelli*) show predator-specific behavior when presented with auditory or visual cues related to their natural predators, which are leopards, eagles and snakes (Lemasson et al., 2009); males behaved conspicuously towards both eagle and leopard models, whereas, females behaved conspicuously towards the leopard but were cryptic to the eagle. Additionally, individuals were found to ascend upon hearing a leopard or after detecting a viper but descend in response to eagle shrieks. Barbary macaques were shown to display some predator-specific behavior, too, as they produced different alarm calls in response to dogs, humans and snakes (Fischer & Hammerschmidt, 2001); and when these alarm calls were played back, their response varied too – startle and escape responses occurred more often after dog alarm calls than calls in response to humans. In contrast, calls related to snakes did not elicit any specific antipredator behavior. Thus, while there may be interindividual differences in the behavioral response to predators due to differences in individual hemispheric specialization, these responses can also vary between different predators.

The current study sets out to identify whether there is a link between lateralization and the behavioral response of two captive groups of Barbary macaques (n = 22) in the context of predation. We hypothesize that the macaques would show individual but not group-level hand preference for unimanual foraging. We furthermore expected that the varying degree of bias in hand preference would relate to differentiated behavioral responses during predator exposure, which we quantified here by looking at focus and tension behaviors after predator exposure. Therefore, we hypothesize a relationship between the direction of hand preference and the intensity of focus and tension behavior. Left-handed individuals are expected to display a higher frequency of focus and tension behavior than right-handed individuals, as the right hemisphere is specialized for predator detection and spontaneous behavioral responses. Also, we predict that this relationship may vary with regard to the two predator models.

## 2. Materials and Methods

### 2.1. Subjects and study sites

We tested twenty-two adult Barbary Macaques (female = 16, male = 8) in this study (see Table S1). The age of macaques ranged from 4 to 20 years (11.1 ± 4.9 years) at the start of data collection. Nine individuals (female = 8, male = 1) were socially housed in a single group along with four infants under a year old at Apenheul Primate Park in Apeldoorn, the Netherlands. Thirteen individuals (female = 8, male = 5) were housed in a social group at Gaia Zoo in Kerkrade, the Netherlands. At Apenheul Primate Park, the Barbary macaques were housed in an outdoor enclosure with a creek, boulders, wooden climbing structures and synthetic rock-like plateaus. The macaques at Gaia Zoo had access to an indoor enclosure and an outdoor one with several trees and climbing structures on a hill-like terrain. At both zoos, the diet consisted of monkey pellets and vegetables and was supplemented with smaller food items such as grains, nuts and seeds. Food was provisioned multiple times a day, and water was available *Ad libitum*.

### 2.2. Data collection and experimental design

#### 2.2.1. Hand-use

We conducted 10-minute continuous focal observations of all the individuals; videos were recorded using a Canon Legria HF R806 camera. The observation sessions typically started after the macaques were given smaller food items such as monkey pellets. We conducted the observations two or three days a week; note that individuals were never observed twice in a row and on the same day. Data were collected from March to May 2022 at Apenheul Primate park and from September to October 2022 at Gaia Zoo. Eight foraging videos were collected on each focal animal except for one individual with seven focal sessions (total observation = 1760 minutes). After excluding time out of sight, the remaining 1714 minutes of data were used for the analyses (average per focal: mean ± standard deviation: 77.9 ± 7.1 minutes). The use of specific hands by the macaques was noted during feeding from the ground (Figure 1b, 1c) while assessing hand preference (see section 2.3.1 for details).

**Figure 1.**
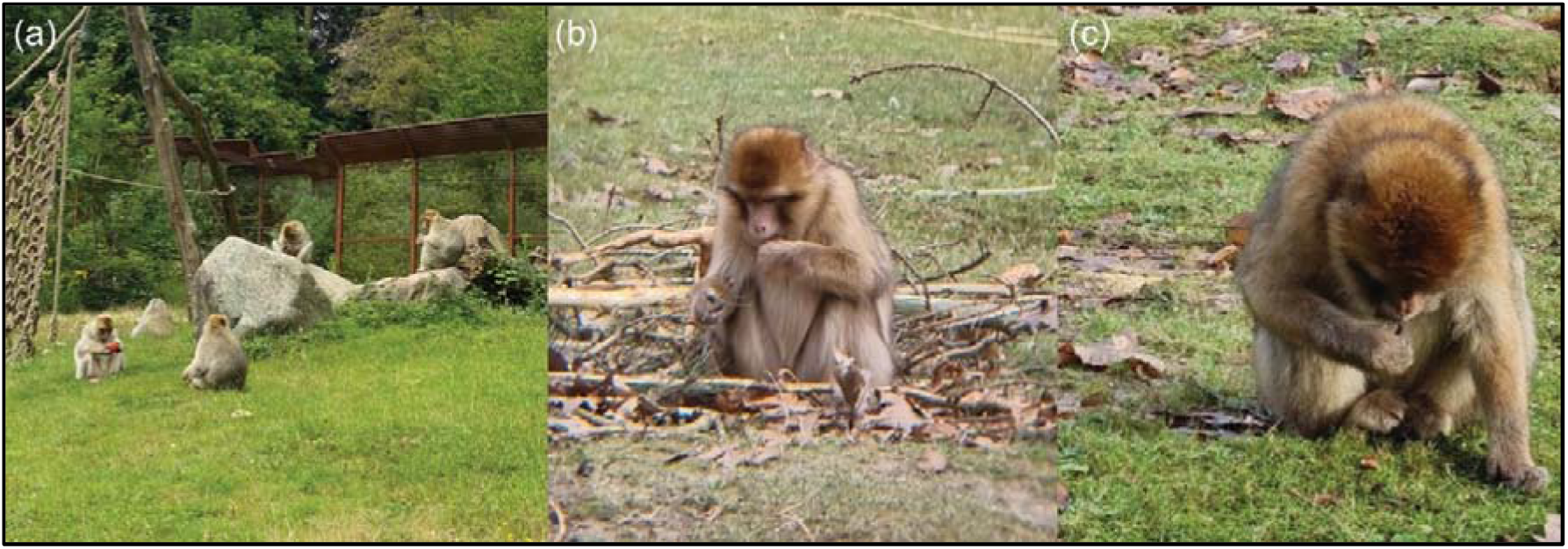
(**a**) Figure showing an example of the foraging behavior of multiple individuals within the group at Gaia Zoo; **(b)** Example of left-hand unimanual foraging; (**c**) Example of right-hand unimanual foraging.

#### 2.2.2. Behavioral response to predators

We conducted experiments to quantify the behavioral responses of the macaques toward potential predator models. A plush large cat/tiger (∼100 cm, Figure S1a) and a rubber snake (∼150 cm, Figure S1b) with markings similar to that of a reticulated python were used (cf. Kluiver et al., 2022). Although the tiger is not a natural predator of Barbary macaques, the cat model might have resembled a leopard to the Barbary macaques (Stein & Hayssen, 2013); besides, the model might also have resembled a large genet, a small predator with a cat-like body, pale fur, black spots and a ringed tail (Larivière & Calzada, 2001; Majolo et al., 2013). The other predator model that was used resembled a reticulated python. Snakes elicit fear in primates, and Barbary macaques are known to respond to numerous species of snakes, even those that pose no serious threat (Isbell, 2006; Roberts et al., 2008). Both predator models were thus ecologically relevant, and both also did trigger antipredator responses by the macaques.

The models were placed on the ground, one meter from the fence, but out of reach of the individuals. We placed the models in such a way that they were facing the macaques. The experiments commenced when a predator model became visible to the macaques either by lifting a blanket or moving it in sight. From uncovering or placing a model, we recorded the behavioral responses of the macaques for 30 minutes, after which the model was removed. The experiments were recorded using a video camera mounted on a tripod.

We repeated each predator exposure session once to obtain enough data on each individual. At Apenheul Primate Park, the first round of predator experiments was conducted in February 2022, when the park was closed to visitors. A second round occurred in May 2022, and experiments were conducted before visitors reached the enclosure. The response to the two different types of models was tested in separate weeks during both rounds. At Gaia Zoo, two rounds were carried out as well, but data from the first round could not be included due to the limited visibility of the macaques. The second round of experiments at Gaia Zoo was conducted at the end of September and beginning of October 2022.

Not all individuals were visible during the predator exposure sessions as they remained in parts of the enclosure out of sight of the camera and potentially out of sight of the predator model. After removing the individuals observed for less than 5 minutes, on average, 22.1 ± 11.7 minutes of data were obtained per focal individual in sight (n = 12) during the tiger model. For the python model, it was 15.8 ± 7.7 minutes (n = 15). We specifically looked at the focus and tension behavior of these individuals during these predator exposures (see section 2.3.2 for details).

### 2.3. Data coding and preparation

The videos were coded in BORIS (v7.13.6) (Friard & Gamba, 2016) by two experimenters (ESJD and EB). In order to calculate the inter-rater reliability (IRR), both experimenters coded 160 minutes of data (translates to 8.6% of all collected data). IRR was found to be very high (ICC (3,k) = 1, p < 0.001).

#### 2.3.1. Hand preference

An extensive ethogram was followed, which consisted of foraging states, events of hand-use during foraging and durational movements (Figure 1, Table 1).

**Table 1.**
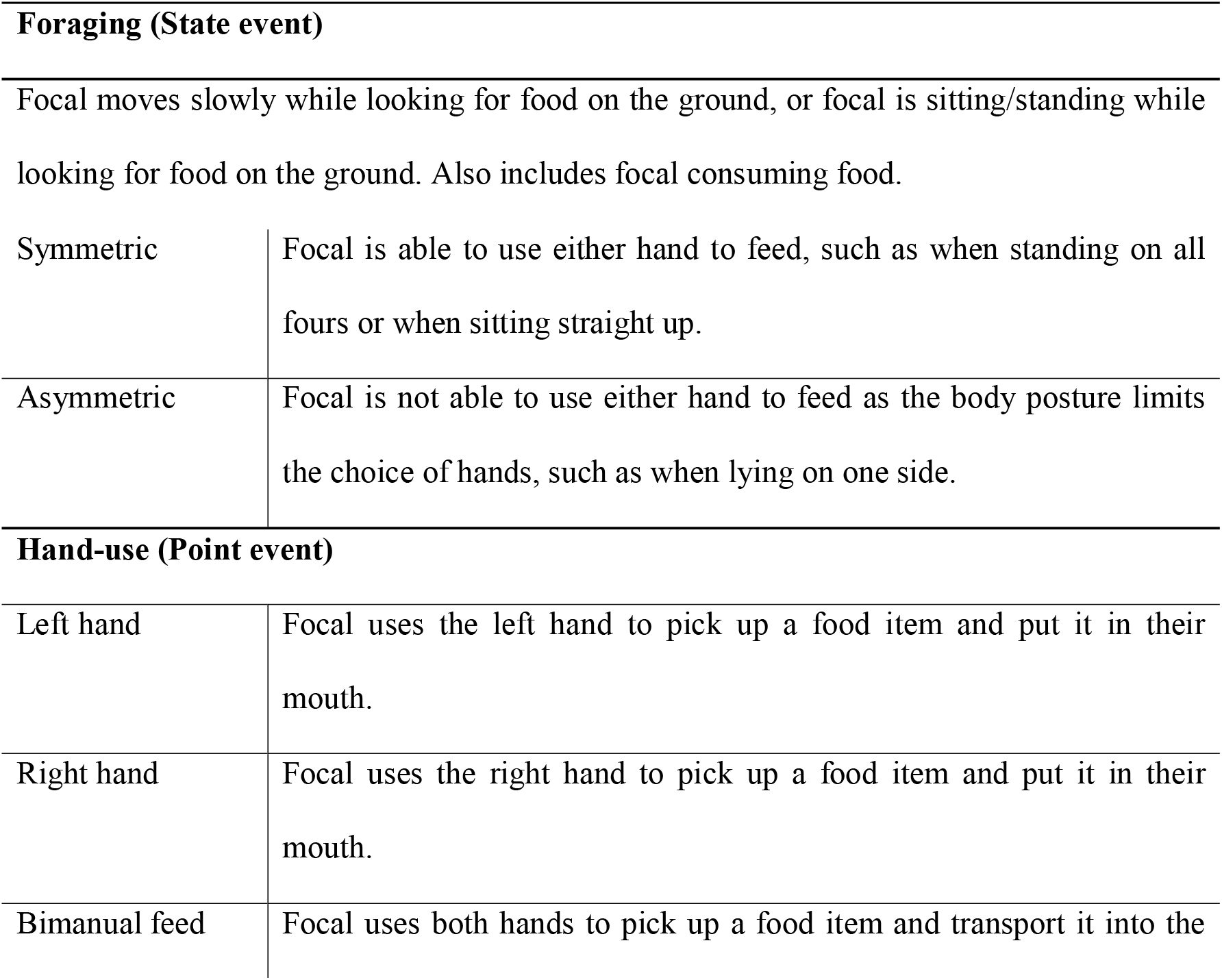
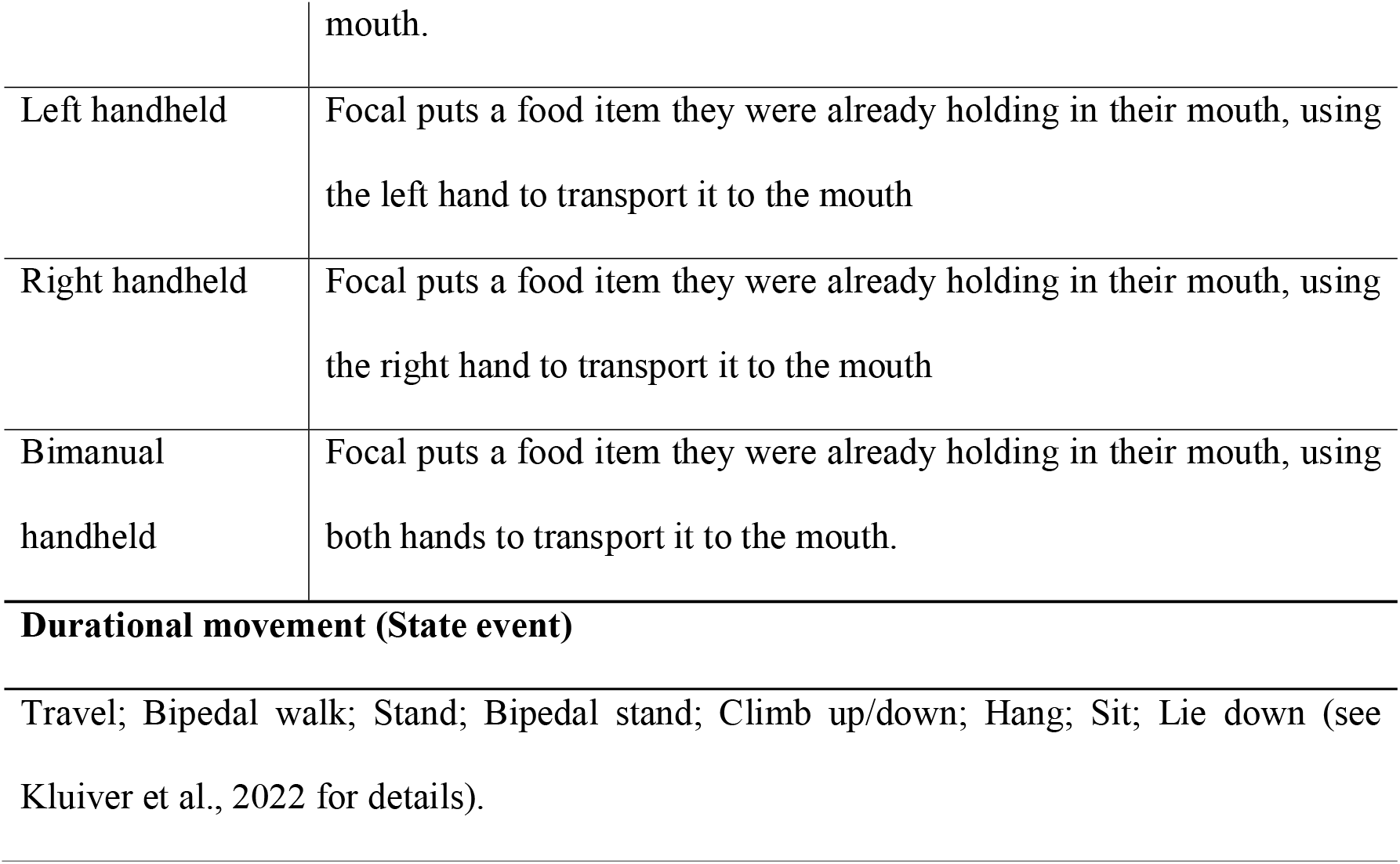
Ethogram of foraging behaviors and their definitions.

For each individual, the total number of instances of feeding with the left hand and feeding with the right hand was calculated, not including handheld and bimanual feeding. In addition to this, the hand-use events were split up into foraging bouts. A new bout was said to start when either the durational movement changed between two instances of hand use or when there was a period (>10 seconds) without foraging between one event and the next (Regaiolli et al., 2018). For each bout, the first occurrence of hand use was determined. The total number of bouts started with a left, or right-hand unimanual feeding was calculated for each focal, omitting bouts started with handheld or bimanual feeding.

As a measure of hand preference, Handedness index (HI) scores were calculated based on the instances and the bouts (Regaiolli et al., 2018; Schmitt et al., 2008). The HI score ranged between -1 and 1, with negative scores relating to a left-side bias and positive scores to a right-side bias. We calculated the HI score for each individual in the following way –

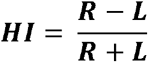

[R is the number of instances or bouts attributed to the right hand, and L is the number attributed to the left hand (Canteloup et al., 2013)]

#### 2.3.2. Behavioral response to predators

The response of individuals to the presence of a predator model was determined by the frequency or occurrence of tension and focus behavior (Table 2). The tension and focus behavior displayed by an individual were combined to create a single measure of the behavioral response as both categories of behavior can be classified as antipredator responses, and there were few occurrences per category, per individual, and per predator type. A total of 797 occurrences of focus and tension behavior were noted. The macaques were not equally visible for the entire length of the experiments; therefore, we corrected them for the time out of sight. We calculated the frequency of these behaviors per minute observed per individual. The response to the tiger model was determined separately from that of the python model.

**Table 2.** Ethogram of tension and focus behavior, and their definitions.

| **Tension** |  |
| --- | --- |
| Avoid conflict | Focal focuses on a conflict without approaching it and moves away in the opposite direction. |
| Body shake | Focal rapidly turns the whole body in at least two different directions. |
| Head turn | While travelling or standing, the focal turns the head in a quick, jerky movement followed by a change of direction. |
| Scratch | Focal uses finger, hand, or foot to rake across own skin. |
| Vigilance | Focal has a tense body posture and looks around with hasty movements of head and/or eyes without an imminent reason. |
| Yawn | Focal opens their mouth wide and inhales intensely, which can be seen by the expansion of the chest. |
| **Focus** |  |
| Attention | Focal looks towards a specific situation, individual or object with a clear focus without moving its head and eyes and with a frozen body posture. |
| Look around | Focal moves its head in at least three different directions without a clear focus. |

### 2.4. Statistical analyses

#### 2.4.1. Hand preference

The preferred hand for unimanual foraging was determined for each individual. If the HI score based on instances had the same sign as the HI score based on bouts, an individual was labelled as left or right-handed, or else they were labelled as ambiguous (Hopkins, 2013). Next, following the recommendations of Hopkins (2013), a possible correlation between the HI score based on instances and the HI score based on bouts was investigated by calculating Pearson’s correlation coefficient. If a strong and positive correlation was found, all further analyses would be based on the handedness index calculated for instances rather than bouts (Hopkins, 2013; Meguerditchian et al., 2010). The possible effect of age and sex on HI was tested using a linear mixed effect model (LMM) with the *lmerTest* package of R (Kuznetsova et al., 2017). We added group as a random effect in the model. Finally, the possibility of a group-level hand preference was investigated using the Kolmogorov-Smirnov test, which reports whether the HI scores differ from a normal distribution (Schmitt et al., 2008).

#### 2.4.2. Hand preference as a predictor of response to predators

We used an LMM to investigate the relationship between hand preference and the behavioral response to predators. Only individuals observed for at least five minutes for a type of predator model were included. The frequency of tension and focus behavior combined was scaled by calculating z-scores. We included this corrected combined frequency of focus and tension behavior as the response variable, HI scores and predator types as fixed effects, sex and age as control variables, and group as a random effect. To examine the explanatory value of our model, we conducted a null versus full model comparison. Additionally, model diagnostics (normality, dispersion, and outlier) were checked using the *DHARMa* package of R (Hartig, 2022). Here, we only report on models that differed significantly from the null model (comparisons and model diagnostics are presented in Supplementary Materials: Table S3 and S4). We performed all analyses in R (version 4.1.2; R Core Team, 2022) using the RStudio interface (version 2022.07.01, Build 554).

## 3. Results

### 3.1. Hand preference

By comparing the HI scores based on bouts and instances of hand use, of the twenty-two individuals, nine were labelled as left-handed, eleven as right-handed, and two were ambiguous for hand preference during unimanual foraging (Figure 2). The HI calculated based on bouts was strongly positively correlated with the HI based on instances (Pearson’s product-moment correlation: r = 0.91, 95% CI [0.79, 0.99], p < 0.001. Therefore, in all further analyses, the HI based on instances rather than bouts were used (cf. Hopkins, 2013; Meguerditchian et al., 2010). All individuals were included in further analyses, including those labelled ambiguous based on their HI for foraging bouts. No group-level hand preference was found for the HI based on instances (Kolmogorov-Smirnov test: D = 0.25, p = 0.11). Using an LMM, the effect of age and sex on HI was tested while controlling for group identity. Age had no impact on the HI (LMM: t(19) = 0.53, p = 0.60), while for sex, a non-significant trend was found (LMM: t(19) = 1.80, p = 0.09) that showed that males might have a higher HI than females (Effect = 0.38 ± 0.21; see Figure 2).

**Figure 2.**
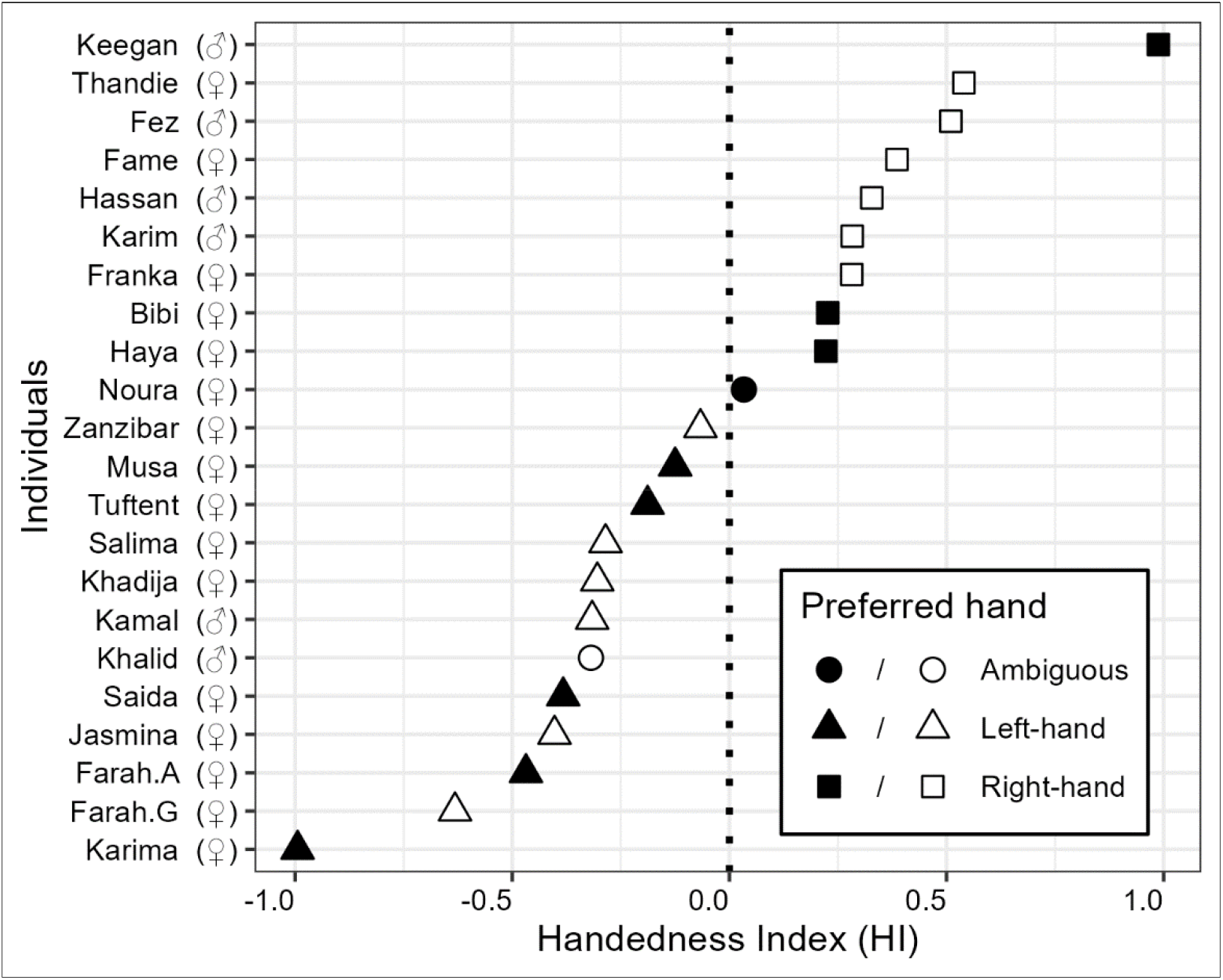
Scatterplot of the handedness index scores of each individual. The different shapes indicate the preferred hand of the individuals based on the labelling criteria. Filled shapes indicate individuals from Apenheul, and outlined shapes from Gaia Zoo. The dotted vertical line indicates HI = 0. To the left of this line, the HI indicates a left-hand bias and to the right, a right-hand bias.

### 3.2. Hand preference and the behavioral response to predators

We found a significant effect of HI on the behavioral responses to predator models (LMM: t = -3.436, p = 0.002; Table S2). Individuals with a stronger left-hand preference elicited more frequent tension and focus behavior than right-handed individuals (Figure 3). Similar to HI, we also found an effect of the predator type (t = -2.796, p = 0.01; Table S2). While the python model did affect individuals, the tiger model particularly had a stronger influence in eliciting tension and focus behavior by the individuals. Furthermore, an interaction effect between HI and the type of predator model was observed (LMM: t = 2.521, p = 0.019; Table S2). An in-depth analysis of the interaction effect revealed that the original main effect in which individuals with a stronger left-hand preference (i.e., HI < 0) elicited more frequent focus and tension behavior, was significantly stronger in response to the tiger than to the python model (t-ratio = 2.631, p = 0.02; Figure 3, Table S2). We did not find any effect of age (t = 1.070, p = 0.29, Table S2). Finally, a non-significant trend was noticed for the effect of sex on the behavioral response to predators (t = 1.918, p = 0.06, Table S2), with males exhibiting more frequent tension and focus behavior than females.

**Figure 3.**
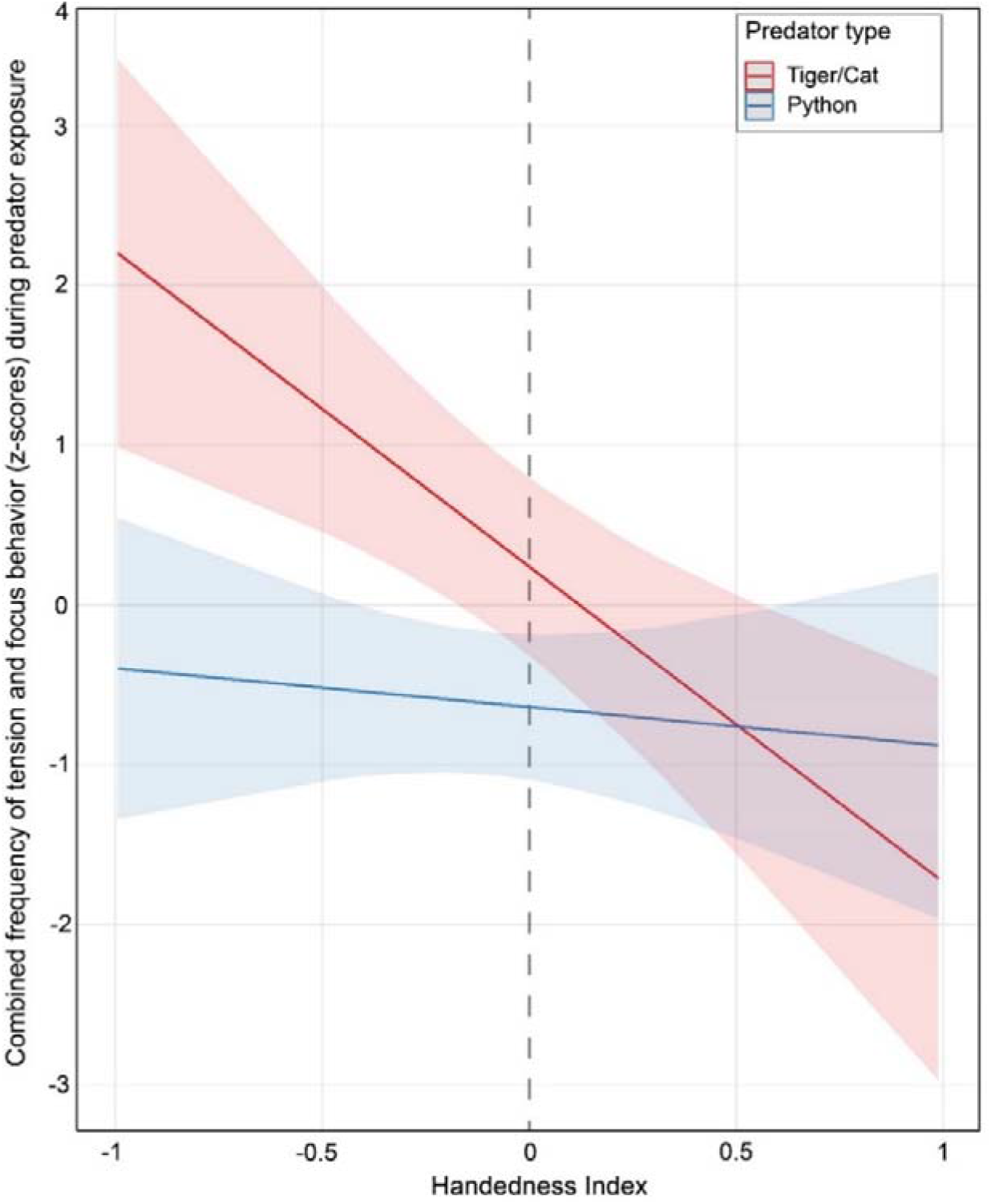
The combined frequency of tension and focus behavior (z-scores) and its relation to an interaction between handedness index and predator type or context. The solid red and blue lines indicate the trend of the two different predator types (red = tiger, blue = python) surrounded by 95% confidence intervals. The left and right sides of the vertical dashed line denote left and right-hand preferences, respectively.

## 4. Discussion

This study examined the relationship between hand preference and behavioral responses of Barbary macaques to potential predation threats. Individual hand preference for unimanual reaching was identified in 20 out of 22 individuals, with no observed group-level bias. Individuals differed in their behavioral responses to potential predators, which was predicted by hand preference, predator type and their interactions. In line with our hypothesis, the hand preference predicted the response to predators; individuals with a stronger left-hand than right-hand bias for unimanual foraging, in general, displayed more frequent tension and focus behavior. The tiger model particularly had a stronger effect on the individuals than the python. Furthermore, an interaction effect between hand preference and predator type implied that the tiger, compared to the python model, had a more substantial influence on individuals with a left-hand than a right-hand bias.

### 4.1. Hand preference

Hand preference during unimanual reaching in Barbary macaques was hypothesized to be an estimator of hemispheric dominance. As such, hand preference for unimanual feeding was expected at an individual but not group level. In line with the predictions, no group-level hand preference was found in this study. Previous studies on Barbary macaque hand preference have also reported the absence of group-level lateralization for unimanual reaching (Baldachini et al., 2021). Here we found evidence of unambiguous hand preference for unimanual foraging in 20 of the 22 individuals tested. This translates to 91% of the sample, while earlier studies on Barbary macaques reported 35%, 47%, and 75% of individuals to be significantly lateralized (Schmitt et al., 2008; Regaiolli et al., 2018; Baldachini et al., 2021). The higher proportion of lateralized individuals reported here compared to earlier research may be attributed to methodological differences rather than differences among the investigated populations. Overall, the presence of individual but not group-level lateralization for unimanual foraging that we observed aligned with the hypothesis that unimanual reaching is an estimator of hemispheric dominance in Barbary macaques.

### 4.2. Predator type and hand preference as predictors of behavioral response

Both the predator models used in this study elicited antipredator behavior, substantiating their ecological relevance as threats. However, the intensity of effects on the individuals varied with the type of predator model; overall, the tiger model elicited a stronger response than the python model. This observation was in line with previous studies that showed varying antipredator responses to different predators (Lemasson et al., 2009). Barbary macaques were found to display antipredator behavior even to play-backs of dog alarm calls, whereas snake alarm calls failed to elicit a response (Fischer & Hammerschmidt, 2001). Similarly, the tiger model may have posed a bigger immediate threat than the python in the current study. Consequently, the tiger model caused a higher frequency of tension and focus behavior and, therefore, a stronger behavioral response by the individuals. These results highlight the importance of behavioral variation with regard to context-specificities, even with a single ecological event of predation.

In addition, independent of the type of predator model, individuals with a stronger left- hand preference displayed a higher frequency of tension and focus behavior than individuals with a stronger right-hand preference. This was in line with the expectation that left-handed individuals would be more reactive to (potential) predators than right-handed individuals, as the right hemisphere is specialized for predator detection and spontaneous behavioral responses (Rogers, 2002, 2010). Finally, a significant interaction effect between handedness index and predator type was observed, where individuals with a left-hand preference showed a more intense reaction to the tiger than the python model. The tiger model was indeed perceived as a greater threat than the python, and coupled with this, hand preference predicted the behavioral responses of Barbary macaques. This suggests an elevated level of reactivity in left-handed individuals while dealing with varying intensities of threat. Although these results indicate that the neural mechanisms may potentially differ when processing predation threats in left and right-handed individuals, it is at the same time challenging to ascertain the specific underlying neural mechanisms non-invasively. Consistent with our findings, previous research on Geoffrey’s marmosets found that left-handed individuals exhibit longer freezing behavior than right-handed individuals in response to hawk calls (Braccini & Caine, 2009). However, right-handed common marmosets are known to produce frequent mobbing calls and perform more head cocking and parallax movements than left-handed marmosets when confronted with a threat (Gordon & Rogers, 2010). This could contradict the pattern found in Geoffrey marmosets, yet Gordon and Rogers (Gordon & Rogers, 2010) argued that these right-handed individuals could be more proactive, as they were explorative towards threats instead of showing signs of withdrawal. Our parameters, i.e., the focus and tension, can be seen as a measure of reactivity, withdrawal or anxiety rather than a proactive investigative response. This is primarily due to the inclusion of behaviors - scratching and yawning, commonly grouped under self-directed behavior and used as indicators of anxiety or stress (Maestripieri et al., 1992; Castles et al., 1999; Castles & Whiten, 1998). As such, the relationship among handedness index, predator type and behavioral response in Barbary macaques can be interpreted as left-handed individuals being more reactive than right-handed individuals, in line with the lateralized brain function and results of earlier studies.

Predator attacks are highly stressful events (Cheney & Seyfarth, 2009), and the response to the presentation of a predator model can thus also be indicative of fear and anxiety (Barros et al., 2008; Carter et al., 2012). While the right hemisphere is dominant for predator detection and spontaneous behavioral responses, it is also involved in the expression of negative emotions such as fear and stress response (Ocklenburg et al., 2016; Rogers, 2002, 2010). Left-handed individuals, considered more reactive than right-handed individuals, could thus be expected to display both stronger behavioral and stress responses to such conditions. However, the relationship between behavioral syndromes, such as reactive and proactive coping styles, which are temporally and contextually consistent correlated behaviors, along with (neuroendocrine) stress reactivity, are hard to disentangle (Koolhaas et al., 2010; Rogers, 2018). Reactive individuals may show the highest hypothalamic–pituitary– adrenal (HPA) axis response, but this varies across species. For example, while there is a positive correlation between right frontal cortex activity and plasma cortisol levels in rhesus macaques (Kalin et al., 1998), a general trend of these levels being lowest in left-handed rhesus macaques is also evident (Westergaard et al., 2003, 2004). Furthermore, for some dimensions, stress reactivity is independent of whether an individual is considered to have a more reactive or proactive behavioral syndrome (Koolhaas et al., 2010).

Research into the potential link between hand preference and stress response has hitherto generated mixed results. In common marmosets, left-handed individuals have more prolonged elevated cortisol levels, possibly indicating that they are more reactive to stress than right-handed individuals (Rogers, 2009). However, a different study reported that the basal cortisol of left-handed common marmosets was lower than that of right-handed individuals and that there is no difference in reactivity between left- and right-handed individuals (Vaughan et al., 2019). In Bonobos (*Pan paniscus*), no relationship between handedness and reactivity was found when investigating self-directed behavior and performance in cognitive tasks (Laméris et al., 2022). Regardless of whether the tension and focus behavior in response to predator models is solely indicative of the behavioral response or an affective one, our findings align with the lateralized brain function, i.e., generated evidence of individuals with a left-hand preference being reactive. Nonetheless, to distinguish the behavioral syndrome of left- and right-handed Barbary macaques from their stress reactivity, it would be valuable to conduct studies using both behavioral and physiological measures.

### 4.3. Potential influence of sex on the behavioral response to predators

A non-significant trend was found, indicating that males may have a higher frequency of tension and focus behavior in response to the predators than females. The behavioral response to predators is sex-specific in several primate species, such as Campbell’s monkeys (Lemasson et al., 2009), white-fronted capuchins (*Cebus albifrons*) and tufted capuchins (*C. apella*) (van Schaik & van Noordwijk, 1989). In general, males show more vigilance and engage in riskier antipredator behavior than females, such as in multiple species of *Cercopithecus* monkeys (Gautier-Hion et al., 1983; Zuberbühler et al., 1997). Thus, the observed trend that suggests sex influenced the behavioral response to predators in Barbary macaques was in line with the general pattern of sexual differences in antipredator behavior in primates. Nevertheless, our study suffered from a low number of males, thus calling for caution during interpretation, and future studies with more males are needed.

### 4.4. Potential shortcomings

Despite testing twice, only one round of predator exposure experiments at Gaia Zoo could be included. As a result of this and other limitations to the data, 27 observations of only 17 different individuals could be included in the final model that related handedness to the predator responses, while the full sample consisted of 22 individuals. It would thus be valuable to repeat this study in a larger sample of (Barbary) macaques. It could also be interesting to investigate the link between sex and response to predators in a larger sample of Barbary macaques since only five males could be included in this study.

Only tension and focus behaviors were used to determine the behavioral response to predators, but different predators may elicit a range of antipredator behavior (Lemasson et al., 2009). Thus, in-depth information on antipredator behavior should be collected in future studies to determine the predictive ability of hand preference.

Finally, we conducted a group-level experiment; therefore, the individual responses might have been influenced by the presence and/or reaction of group members. Left-handed individuals are suggested to be less responsive to and of influence on their social group than right-handed individuals (Gordon & Rogers, 2010). Nonetheless, a group-level experiment best resembles how predators are detected and responded to under wild or semi-wild circumstances, and as such, also mimics best the actual selection pressures during evolution. Therefore, this was an ecologically valid context to investigate the effect of lateralization of hand preference on individual differences in behavior.

## 5. Conclusions

Hand preference was a reliable predictor of the frequency of tension and focus behavior of Barbary macaques in the context of predation. The extent of left-hand preference was positively related to the frequency of focus and tension behaviors. The direction of this effect was in line with the right-hemispheric specialization for predator detection, spontaneous behavioral responses and negative emotions. As such, our results were consistent with the lateralized brain function hypothesis and suggested that cerebral lateralization could be one of the neural mechanisms underlying individual differences in context-specific bevaiours. Furthermore, an interaction between hand preference and the predator context translates to (i) a higher reactivity of left-than right-handed individuals and (ii) potentially different neural information processing mechanisms. As predator exposure is just one of many ecologically relevant contexts, lateralization may also result in inter-individual variation in other context-specific behaviors, both in Barbary macaques and other non-human primates. Therefore, future research on other non-human primates and other ecologically relevant contexts building onto the current framework could reveal clues about the evolution of brain lateralization and potentially related behavioral predispositions.

## Supplementary Materials

Supplementary file S1: Datasets.

Supplementary file S2: R-scripts.

**Table S1**. Description of study animals, including individual ID, housing location, year of birth and sex.

**Figure S1**. (a) Tiger/cat model used for predator exposure experiment. (b) Python model used for predator exposure experiment.

**Table S2**. Test statistics for the fixed effects of a mixed effect model testing the influence of handedness, context (predator type), sex and age on the corrected frequency of tension and focus behaviors.

**Table S3**. Model (full mixed effect model testing the influence of handedness, context (predator type), sex and age on the corrected frequency of tension and focus behaviors.) diagnostics reporting on the (a) uniformity, (b) dispersion, (c) outliers of the residuals.

**Table S4**. Test statistics for the null model versus full model comparison.

## Author Contributions

Conceptualization: ESJD and DB; methodology: ESJD, DB and JJMM; formal analysis: ESJD, EB and DB; investigation: ESJD and EB; resources: DB and JJMM; data curation: ESJD; writing: ESJD; writing—review and editing: DB and JJMM; supervision: DB and JJMM; funding acquisition: DB and JJMM. All authors have read and agreed to the published version of the manuscript.

## Institutional Review Board Statement

This study was non-invasive, requiring no ethical approval according to the European Directive 2010/63. Permission to conduct research was obtained from the Apenheul Primate Park and Gaia Zoo, and signed letters have been provided. The experimenters followed all internal protocols and guidelines of the zoos while working with the non-human primates.

## Data Availability Statement

The data presented in the study are available in the supplementary files. Supplementary File S1: Datasets. Supplementary File S2: R-scripts.

## Acknowledgements

We thank Lisette van den Berg for welcoming us to the Apenheul primate park. We are thankful to Emile F. Prins and Thomas Remmits for allowing and supporting the studies at Gaia zoo. We sincerely thank all the caretakers and zookeepers from the Apenheul and Gaia zoos for their help during the study.

## References

Anderson, C. M. (1986). Predation and primate evolution. Primates, 27(1), 15–39. https://doi.org/10.1007/BF02382520

Baldachini, M., Regaiolli, B., Llorente, M., Riba, D., & Spiezio, C. (2021). The Influence of Target Animacy and Social Rank on Hand Preference in Barbary Macaques (Macaca sylvanus). International Journal of Primatology, 42(2), 155–170. https://doi.org/10.1007/s10764-020-00193-0

Barros, M., Maior, R. S., Huston, J. P., & Tomaz, C. (2008). Predatory Stress as an Experimental Strategy to Measure Fear and Anxiety-related Behaviors in Non-human Primates. Reviews in the Neurosciences, 19(2–3), 157–170. https://doi.org/10.1515/REVNEURO.2008.19.2-3.157

Bautista, J. (2019). Golden Eagles (Aquila chrysaetos) as potential predators of Barbary Macaques (Macaca sylvanus) in northern Morocco: Evidences of predation. 15, 181–188.

Bisazza, A. J. Rogers, L., & Vallortigara, G. (1998). The Origins of Cerebral Asymmetry: A Review of Evidence of Behavioural and Brain Lateralization in Fishes, Reptiles and Amphibians. Neuroscience & Biobehavioral Reviews, 22(3), 411–426. https://doi.org/10.1016/S0149-7634(97)00050-X

Braccini, S. N., & Caine, N. G. (2009). Hand preference predicts reactions to novel foods and predators in marmosets (Callithrix geoffroyi). Journal of Comparative Psychology, 123(1), 18.

Cameron, R., & Rogers, L. J. (1999). Hand preference of the common marmoset (Callithrix jacchus): Problem solving and responses in a novel setting. Journal of Comparative Psychology, 113(2), 149–157. https://doi.org/10.1037/0735-7036.113.2.149

Canteloup, C., Vauclair, J., & Meunier, H. (2013). Hand preferences on unimanual and bimanual tasks in Tonkean macaques (Macaca tonkeana). American Journal of Physical Anthropology, 152(3), 315–321. https://doi.org/10.1002/ajpa.22342

Carter, A. J., Marshall, H. H., Heinsohn, R., & Cowlishaw, G. (2012). How not to measure boldness: Novel object and antipredator responses are not the same in wild baboons. Animal Behaviour, 84(3), 603–609. https://doi.org/10.1016/j.anbehav.2012.06.015

Castles, D. L., & Whiten, A. (1998). Post-conflict Behaviour of Wild Olive Baboons. II. Stress and Self-directed Behaviour. Ethology, 104(2), 148–160. https://doi.org/10.1111/j.1439-0310.1998.tb00058.x

Castles, D. L., Whiten, A., & Aureli, F. (1999). Social anxiety, relationships and self-directed behaviour among wild female olive baboons. Animal Behaviour, 58(6), 1207–1215. https://doi.org/10.1006/anbe.1999.1250

Caspar, K. R., Pallasdies, F., Mader, L., Sartorelli, H., & Begall, S. (2022). The evolution and biological correlates of hand preferences in anthropoid primates. Elife, 11, e77875. https://doi.org/10.7554/eLife.77875

Cheney, D. L., & Seyfarth, R. M. (2009). Chapter 1 Stress and Coping Mechanisms in Female Primates. In Advances in the Study of Behavior (Vol. 39, pp. 1–44). Academic Press. https://doi.org/10.1016/S0065-3454(09)39001-4

Désiré, L., Boissy, A., & Veissier, I. (2002). Emotions in farm animals: A new approach to animal welfare in applied ethology. Behavioural Processes, 60(2), 165–180. https://doi.org/10.1016/S0376-6357(02)00081-5

Fernández-Lázaro, G., Latorre, R., Alonso-García, E., & Barja Núñez, I. (2019). Nonhuman primate welfare: Can there be a relationship between personality, lateralization and physiological indicators? Behavioural Processes, 166, 103897. https://doi.org/10.1016/j.beproc.2019.103897

Fischer, J., & Hammerschmidt, K. (2001). Functional referents and acoustic similarity revisited: The case of Barbary macaque alarm calls. Animal Cognition, 4(1), 29–35. https://doi.org/10.1007/s100710100093

Fischer, J., & Hammerschmidt, K. (2002). An Overview of the Barbary Macaque, Macaca sylvanus, Vocal Repertoire. Folia Primatologica, 73(1), 32–45. https://doi.org/10.1159/000060417

Fooden, J. (2007). Systematic Review of the Barbary Macaque, Macaca Sylvanus (Linnaeus, 1758). Fieldiana Zoology, 2007(113), 1–60. https://doi.org/10.3158/0015-0754(2007)113[1:SROTBM]20.CO;2

Friard, O., & Gamba, M. (2016). BORIS: A free, versatile open-source event-logging software for video/audio coding and live observations. Methods in Ecology and Evolution, 7(11), 1325–1330. https://doi.org/10.1111/2041-210X.12584

Fu, W., Xu, J., Wang, X., Li, Y., He, S., Wang, C., Ren, Y., Yang, B., Wu, T., Wang, Y., & Li, B. (2022). Consistency of limb preference across unimanual feeding, bipedal locomotion, and social grooming in golden snub-nosed monkeys (Rhinopithecus roxellana). Laterality, 0(0), 1–16. https://doi.org/10.1080/1357650X.2022.2141251

Gautier-Hion, A., Quris, R., & Gautier, J.-P. (1983). Monospecific vs polyspecific life: A comparative study of foraging and antipredatory tactics in a community of Cercopithecus monkeys. Behavioral Ecology and Sociobiology, 12(4), 325–335. https://doi.org/10.1007/BF00302901

Gordon, D. J., & Rogers, L. J. (2010). Differences in social and vocal behavior between left- and right-handed common marmosets (Callithrix jacchus). Journal of Comparative Psychology, 124(4), 402–411. https://doi.org/10.1037/a0019736

Gordon, D. J., & Rogers, L. J. (2015). Cognitive bias, hand preference and welfare of common marmosets. Behavioural Brain Research, 287, 100–108. https://doi.org/10.1016/j.bbr.2015.03.037

Hartig, F. (2022). DHARMa: Residual Diagnostics for Hierarchical (Multi-Level / Mixed) Regression Models. https://CRAN.R-project.org/package=DHARMa

Hook, M. A., & Rogers, L. J. (2000). Development of hand preferences in marmosets (Callithrix jacchus) and effects of aging. Journal of Comparative Psychology, 114(3), 263.

Hopkins, W. D. (2013). Independence of data points in the measurement of hand preferences in primates: Statistical problem or urban myth? American Journal of Physical Anthropology, 151(1), 151–157. https://doi.org/10.1002/ajpa.22248

Hopkins, W. D., & Bennett, A. J. (1994). Handedness and approach-avoidance behavior in chimpanzees (Pan). Journal of Experimental Psychology: Animal Behavior Processes, 20(4), 413–418. https://doi.org/10.1037/0097-7403.20.4.413

Isbell, L. A. (2006). Snakes as agents of evolutionary change in primate brains. Journal of Human Evolution, 51(1), 1–35. https://doi.org/10.1016/j.jhevol.2005.12.012

Kalin, N. H., Larson, C., Shelton, S. E., & Davidson, R. J. (1998). Asymmetric frontal brain activity, cortisol, and behavior associated with fearful temperament in rhesus monkeys. Behavioral Neuroscience, 112(2), 286–292. https://doi.org/10.1037/0735-7044.112.2.286

Kappeler, P. M., & van Schaik, C. P. (2002). Evolution of Primate Social Systems. International Journal of Primatology, 23(4), 707–740. https://doi.org/10.1023/A:1015520830318

Kluiver, C. E., de Jong, J. A., Massen, J. J. M., & Bhattacharjee, D. (2022). Personality as a Predictor of Time-Activity Budget in Lion-Tailed Macaques (Macaca silenus). Animals, 12(12), Article 12. https://doi.org/10.3390/ani12121495

Koolhaas, J. M., de Boer, S. F., Coppens, C. M., & Buwalda, B. (2010). Neuroendocrinology of coping styles: Towards understanding the biology of individual variation. Frontiers in Neuroendocrinology, 31(3), 307–321. https://doi.org/10.1016/j.yfrne.2010.04.001

Kuběnová, B., Lhota, S., Tomanová, V., Blažek, V., & Konečná, M. (2022). Lion-tailed macaques show a stable direction and reinforcement of hand preference in simple reaching tasks over several years. Journal of Vertebrate Biology, 71(21076), 21076.1–13. https://doi.org/10.25225/jvb.21076

Kuznetsova, A., Brockhoff, P. B., & Christensen, R. H. B. (2017). lmerTest Package: Tests in Linear Mixed Effects Models. Journal of Statistical Software, 82(13), 1–26. https://doi.org/10.18637/jss.v082.i13

Laméris, D. W., Verspeek, J., Salas, M., Staes, N., Torfs, J. R. R., Eens, M., & Stevens, J. M. G. (2022). Evaluating Self-Directed Behaviours and Their Association with Emotional Arousal across Two Cognitive Tasks in Bonobos (Pan paniscus). Animals, 12(21), Article 21. https://doi.org/10.3390/ani12213002

Larivière, S., & Calzada, J. (2001). Genetta genetta. Mammalian Species, 2001(680), 1–6. https://doi.org/10.1644/1545-1410(2001)680<0001:GG>2.0.CO;2

Leliveld, L. M. C., Langbein, J., & Puppe, B. (2013). The emergence of emotional lateralization: Evidence in non-human vertebrates and implications for farm animals. Applied Animal Behaviour Science, 145(1), 1–14. https://doi.org/10.1016/j.applanim.2013.02.002

Lemasson, A., Ouattara, K., & Zuberbühler, K. (2009). Anti-predator strategies of free-ranging Campbell’s monkeys. Behaviour, 146(12), 1687–1708. https://doi.org/10.1163/000579509X12469533725585

MacNeilage, P. F., Rogers, L. J., & Vallortigara, G. (2009). Origins of the left & right brain. Scientific American, 301(1), 60–67.

Maestripieri, D., Schino, G., Aureli, F., & Troisi, A. (1992). A modest proposal: Displacement activities as an indicator of emotions in primates. Animal Behaviour, 44(5), 967–979. https://doi.org/10.1016/S0003-3472(05)80592-5

Majolo, B., McFarland, R., Young, C., & Qarro, M. (2013). The Effect of Climatic Factors on the Activity Budgets of Barbary Macaques (Macaca sylvanus). International Journal of Primatology, 34(3), 500–514. https://doi.org/10.1007/s10764-013-9678-8

Masilkova, M., Šlipogor, V., Lima Marques Silva, G. H., Hadová, M., Lhota, S., Bugnyar, T., & Konečná, M. (2022). Age, but not hand preference, is related to personality traits in common marmosets (Callithrix jacchus). Royal Society Open Science, 9(10), 220797. https://doi.org/10.1098/rsos.220797

Meguerditchian, A., Calcutt, S. E., Lonsdorf, E. V., Ross, S. R., & Hopkins, W. D. (2010). Brief communication: Captive gorillas are right-handed for bimanual feeding. American Journal of Physical Anthropology, 141(4), 638–645. https://doi.org/10.1002/ajpa.21244

Meguerditchian, A., Vauclair, J., & Hopkins, W. D. (2013). On the origins of human handedness and language: A comparative review of hand preferences for bimanual coordinated actions and gestural communication in nonhuman primates. Developmental Psychobiology, 55(6), 637–650. https://doi.org/10.1002/dev.21150

Nelson, E. L. (2022). Insights Into Human and Nonhuman Primate Handedness From Measuring Both Hands. Current Directions in Psychological Science, 31(2), 154–161. https://doi.org/10.1177/09637214211062876

Ocklenburg, S., Korte, S. M., Peterburs, J., Wolf, O. T., & Güntürkün, O. (2016). Stress and laterality—The comparative perspective. Physiology and Behavior, 164, 321–329. Scopus. https://doi.org/10.1016/j.physbeh.2016.06.020

Öhman, A., & Mineka, S. (2003). The Malicious Serpent: Snakes as a Prototypical Stimulus for an Evolved Module of Fear. Current Directions in Psychological Science, 12(1), 5–9. https://doi.org/10.1111/1467-8721.01211

R Core Team (2022). R: A language and environment for statistical computing. R Foundation for Statistical Computing, Vienna, Austria. https://www.R-project.org/

Raymond, M., & Pontier, D. (2004). Is there geographical variation in human handedness? Laterality, 9(1), 35–51. https://doi.org/10.1080/13576500244000274

Regaiolli, B., Spiezio, C., & Hopkins, W. D. (2018). Hand preference on unimanual and bimanual tasks in Barbary macaques (Macaca sylvanus). American Journal of Primatology, 80(3), e22745. https://doi.org/10.1002/ajp.22745

Roberts, S., McComb, K., & Ruffman, T. (2008). An Experimental Investigation of Referential Looking in Free-Ranging Barbary Macaques (Macaca sylvanus). Journal of Comparative Psychology (Washington, D.C.LJ: 1983), 122, 94–99. https://doi.org/10.1037/0735-7036.122.1.94

Rogers, L. J. (2002). Lateralization in vertebrates: Its early evolution, general pattern, and development. In Advances in the Study of Behavior (Vol. 31, pp. 107–161). Elsevier. https://doi.org/10.1016/S0065-3454(02)80007-9

Rogers, L. J. (2009). Hand and paw preferences in relation to the lateralized brain. Philosophical Transactions of the Royal Society B: Biological Sciences, 364(1519), 943–954. https://doi.org/10.1098/rstb.2008.0225

Rogers, L. J. (2010). Relevance of brain and behavioural lateralization to animal welfare. Applied Animal Behaviour Science, 127(1), 1–11. https://doi.org/10.1016/j.applanim.2010.06.008

Rogers, L. J. (2011). Does brain lateralization have practical implications for improving animal welfare? CAB Reviews: Perspectives in Agriculture, Veterinary Science, Nutrition and Natural Resources, 6(036). https://doi.org/10.1079/PAVSNNR20116036

Rogers, L. J. (2018). Chapter 4—Manual bias, behavior, and cognition in common marmosets and other primates. In G. S. Forrester, W. D. Hopkins, K. Hudry, & A. Lindell (Eds.), Progress in Brain Research (Vol. 238, pp. 91–113). Elsevier. https://doi.org/10.1016/bs.pbr.2018.06.004

Rogers, L. J., & Andrew, R. (2002). Comparative Vertebrate Lateralization. Cambridge University Press. http://ebookcentral.proquest.com/lib/uunl/detail.action?docID=217820

Schmitt, V., Melchisedech, S., Hammerschmidt, K., & Fischer, J. (2008). Hand preferences in Barbary macaques (Macaca sylvanus). Laterality, 13(2), 143–157. https://doi.org/10.1080/13576500701757532

Šlipogor, V., Gunhold-de Oliveira, T., Tadić, Z., Massen, J. J. M., & Bugnyar, T. (2016). Consistent inter-individual differences in common marmosets (Callithrix jacchus) in Boldness-Shyness, Stress-Activity, and Exploration-Avoidance. American Journal of Primatology, 78(9), 961–973. https://doi.org/10.1002/ajp.22566

Soto, C., Gázquez, J. M. M., & Llorente, M. (2022). Hand preferences in coordinated bimanual tasks in non-human primates: A systematic review and meta-analysis. Neuroscience & Biobehavioral Reviews, 141, 104822. https://doi.org/10.1016/j.neubiorev.2022.104822

Stanford, C. B. (2002). Avoiding Predators: Expectations and Evidence in Primate Antipredator Behavior. International Journal of Primatology, 23(4), 741–757. https://doi.org/10.1023/A:1015572814388

Stein, A. B., & Hayssen, V. (2013). Panthera pardus (Carnivora: Felidae). Mammalian Species, 45(900), 30–48. https://doi.org/10.1644/900.1

Thierry, B., Iwaniuk, A. N., & Pellis, S. M. (2000). The Influence of Phylogeny on the Social Behaviour of Macaques (Primates: Cercopithecidae, genus Macaca). Ethology, 106(8), 713–728. https://doi.org/10.1046/j.1439-0310.2000.00583.x

Tomaz, C., Verburg, M. S., Boere, V., Pianta, T. F., & Belo, M. (2003). Evidence of hemispheric specialization in marmosets (Callithrix penicillata) using tympanic membrane thermometry. Brazilian Journal of Medical and Biological Research, 36, 913–918. https://doi.org/10.1590/S0100-879X2003000700012

van Schaik, C. P., & van Noordwijk, M. A. (1989). The special role of male Cebus monkeys in predation avoidance and its effect on group composition. Behavioral Ecology and Sociobiology, 24(5), 265–276. https://doi.org/10.1007/BF00290902

Vaughan, E., Le, A., Casey, M., Workman, K. P., & Lacreuse, A. (2019). Baseline cortisol levels and social behavior differ as a function of handedness in marmosets (Callithrix jacchus). American Journal of Primatology, 81(9), e23057. https://doi.org/10.1002/ajp.23057

Wang, L., Luo, Y., Lin, H., Xu, N., Gu, Y., Bu, H., Bai, Y., & Li, Z. (2022). Performance on inhibitory tasks does not relate to handedness in several small groups of Callitrichids. Animal Cognition. https://doi.org/10.1007/s10071-022-01682-w

Waterman, J. O., Campbell, L. a. D., Maréchal, L., Pilot, M., & Majolo, B. (2020). Effect of human activity on habitat selection in the endangered Barbary macaque. Animal Conservation, 23(4), 373–385. https://doi.org/10.1111/acv.12543

Westergaard, G. C., Chavanne, T. J., Houser, L., Cleveland, A., Snoy, P. J., Suomi, S. J., & Higley, J. D. (2004). Biobehavioural correlates of hand preference in freeLranging female primates. Laterality, 9(3), 267–285. https://doi.org/10.1080/13576500342000086a

Westergaard, G. C., Chavanne, T. J., Lussier, I. D., Houser, L., Cleveland, A., Suomi, S. J., & Higley, J. D. (2003). Left-handedness is Correlated with CSF Monoamine Metabolite and Plasma Cortisol Concentrations, and with Impaired Sociality, in Free-ranging Adult Male Rhesus Macaques (Macaca mulatta). Laterality, 8(2), 169–187. https://doi.org/10.1080/713754484

Wright, L., Watt, S., & Hardie, S. M. (2013). Influences of lateral preference and personality on behaviour towards a manual sorting task. Personality and Individual Differences, 54(8), 903–907. https://doi.org/10.1016/j.paid.2013.01.005

Zonato, A., Gagliardo, A., Bandoli, F., & Palagi, E. (2022). Reaching versus catching: Flexible hand preference in ring-tailed lemurs. Ethology Ecology & Evolution, 0(0), 1–22. https://doi.org/10.1080/03949370.2022.2098382

Zuberbühler, K., Noë, R., & Seyfarth, R. M. (1997). Diana monkey long-distance calls: Messages for conspecifics and predators. Animal Behaviour, 53(3), 589–604. https://doi.org/10.1006/anbe.1996.0334

