## Supplementary Material for "Hand preference predicts behavioral responses to threats in Barbary macaques"

**Table of contents	Page**

**-----------------------------------------------------------------------------------------------------------------**

**Table S1 –** Description of study animals	2

**Figure S1 –** Predator models used in the study	3

**Table S2 –** Effect of Handedness index and predator type on behavior	3

**Table S3 –** Model diagnostics	4

**Table S4 –** Null and Full model comparison	4

**Table S1.** Description of study animals including individual ID, housing location, year of birth and sex.

| ID | Sex | Year of Birth | Location |
| --- | --- | --- | --- |
| Bibi | F | 2014 | Apenheul |
| Farah A | F | 2014 | Apenheul |
| Haya | F | 2016 | Apenheul |
| Karima | F | 2012 | Apenheul |
| Keegan | M | 2009 | Apenheul |
| Musa | F | 2017 | Apenheul |
| Noura | F | 2006 | Apenheul |
| Saida | F | 2001 | Apenheul |
| Tuftent | F | 2017 | Apenheul |
| Farah G | F | 2002 | GaiaZoo |
| Thandie | F | 2010 | GaiaZoo |
| Kamal | M | 2012 | GaiaZoo |
| Khalid | M | 2013 | GaiaZoo |
| Khadija | F | 2014 | GaiaZoo |
| Hassan | M | 2007 | GaiaZoo |
| Franka | F | 2003 | GaiaZoo |
| Zanzibar | F | 2011 | GaiaZoo |
| Jasmina | F | 2012 | GaiaZoo |
| Fame | F | 2004 | GaiaZoo |
| Fez | M | 2011 | GaiaZoo |
| Karim | M | 2012 | GaiaZoo |
| Salima | F | 2013 | GaiaZoo |

**Figure S1.** **(a)** Tiger/cat model used for predator exposure experiment. **(b)** Python model used for predator exposure experiment.

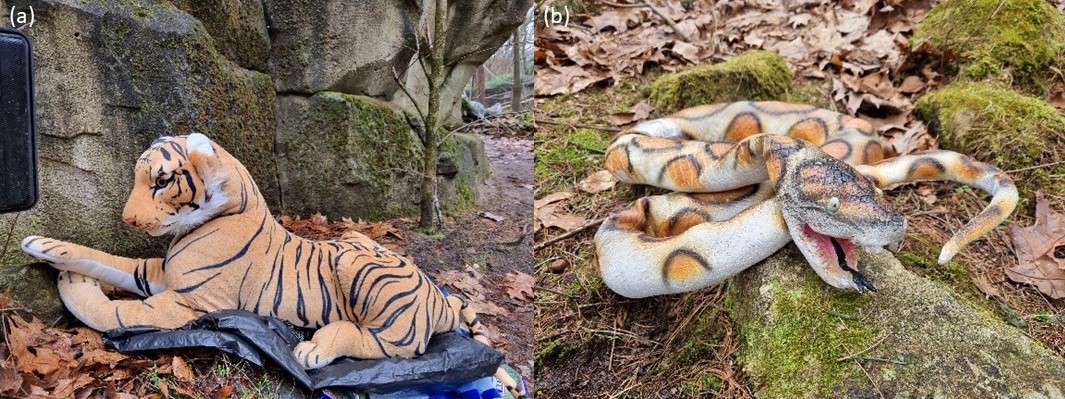

**Table S2.** Test statistics for the fixed effects of a mixed effect model (Model 1) testing the influence of handedness, context (predator type), sex and age on the corrected frequency of tension and focus behaviors.

*Model 1:* Frequency of tension and focus ~ HI * Predator type + Age + Sex + (1|Group/Subject)

| **Measure** | **Estimate** | **Standard error** | **t-value** | **p-value** |
| --- | --- | --- | --- | --- |
| Intercept | -0.3264 | 0.4979 | -0.656 | 0.5192 |
| Handedness index (HI) | -1.9444 | 0.5658 | -3.436 | 0.0024 ** |
| Predator type (Python) | -0.8679 | 0.3104 | -2.796 | 0.0108 * |
| Age | 0.0621 | 0.0581 | 1.070 | 0.2969 |
| Sex (Male) | 0.7220 | 0.3764 | 1.918 | 0.0687 |
| HI * Predator type (Python) | 1.7106 | 0.6785 | 2.521 | 0.0198 * |

Signifiance codes: 0.001 ‘**’ , 0.01 ‘*’.

*Investigation of Interaction Effects:*

Estimate	SE	t-ratio	p-value

Contrasts: HI * Predator (Tiger) – HI * Predator (Python) 0.844	0.321	2.631	0.0220*

**Table S3.** Model 1 diagnostics reporting on the uniformity of the residuals of the full mixed effect model testing the influence of handedness, context (predator type), sex and age on the corrected frequency of tension and focus behaviors.

*(a) Uniformity*:

| Uniformity | |
| --- | --- |
| Test | Asymptotic one-sample Kolmogorov-Smirnov test |
| Data | Scaled residuals of simulation output |
| Result | D = 0.13778, p-value = 0.6846 |
| Alternative | Two-sided |

*(b) Dispersion*:

| Dispersion | |
| --- | --- |
| Test | DHARMa nonparametric dispersion test via sd of residuals fitted vs. simulated |
| Data | Simulation output |
| Result | Dispersion = 0.79653, p-value = 0.512 |
| Alternative | Two-sided |

*(c) Outliers*:

| Outliers | |
| --- | --- |
| Test | DHARMa outlier test based on exact binomial test with approximate expectations |
| Data | Simulation output |
| Results | Outliers at both margin(s) = 0, observations = 27, p-value = 1  95 percent confidence interval: 0.0000000, 0.1277029  Sample estimates: frequency of outliers (expected: 0.007968) 0 |
| Alternative | True probability of success is not equal to 0.00796812 |

**Table S4.** Test statistics for the null model versus full model (Model 1) comparison, performed using ANOVA. Full model is a mixed effects model testing the influence of handedness, context (predator type), sex and age on the corrected frequency of tension and focus behavior while correcting for individual identity nested in group. Null model tests the effect of the random effect of individual identity nested in group in a mixed effect model.

| **Model** | **AIC** | **BIC** | **logLik** | **Deviance** | **Chi2** | **p-value** |
| --- | --- | --- | --- | --- | --- | --- |
| Null | 81.604 | 85.491 | -37.802 | 75.604 |  |  |
| Full | 71.415 | 81.782 | -27.708 | 55.415 | 20.188 | 0.0011 ** |

Significance code: 0.001 ‘**’.

*-----------------------------------------------------------------------------------------------------------------End of Supplementary materials*
